# Red harvester ant (Order: Hymenoptera) preference for cover crop seeds in South Texas

**DOI:** 10.1101/2022.01.14.476276

**Authors:** Lilly Elliott, Daniella Rivera, Adrian Noval, Robin A. Choudhury, Hannah J. Penn

## Abstract

Harvester ants are known to selectively forage seeds, potentially impacting nearby plant community composition. In agricultural areas, harvester ants may be viewed as pests by foraging on crop seeds or as beneficials by preferentially removing weed seeds. However, little work has been done on harvester ant preferences for cover crop seeds. Local observations suggest that ants may take cover crop seeds, but no studies have evaluated ant agricultural impacts or seed preferences in the Lower Rio Grande Valley (LRGV). We examined red harvester ant (*Pogonomyrmex barbatus* Smith) preferences for commonly used cover crop seeds in the LRGV (vetch, oat, fescue, sunn hemp, and radish with wheatgrass as a control) and a commonly used bacterial seed inoculation treatment meant to increase root nodulation. We tested seed sets using choice tests housed in seed depots located within the foraging range of ant colonies with no prior exposure to the selected seeds. Of the evaluated cover crop seeds, wheatgrass and oat were the first to be removed entirely from the depot, with vetch remaining after 24 h. When we inoculated the two most preferred seeds to determine if there was a preference for non-inoculated seeds, we found no difference between inoculated and non-inoculated seeds. There were also significant changes in activity over time for both trials. These data indicate that harvester ant foraging preferences and activity can inform grower management recommendations regarding the risks of using certain cover crops and months sowing should be conducted in fields with known harvester ant presence.

## Introduction

Harvester ants in the genus *Pogonomyrmex* commonly reside in arid to semi-arid regions of the Americas and can be found in a range of habitats including agricultural and peri-urban matrices (Luna et al. 2018; Viera-Neto et al. 2016; Tizón et al. 2010; MacMahon 2000). The state of Texas has 12 species of harvester ants with the red harvester ant (*Pogonomyrmex barbatus* Smith) being the most common in the Lower Rio Grande Valley (LRGV), an agriculturally rich location with a semi-arid sub-tropical climate (Martinez et. al., 2020, Davis, 2016). Harvester ant foraging occurs mainly along trails that extend from the colony to neighboring food sources within their foraging range (Taber, 1999; Traniello, 1989). While foraging trails average 10 m long, colony-dense areas (which may have over 80 nests/hectare) have trails extending up to 60 m from the nest site (Reed and Landolt, 2019). Harvester ants, primarily granivores, use these trails to collect seeds located on the soil surface, often from or surrounding the parent plant (MacKay and MacKay 2002; Taber, 1999). *P. barbatus*, *P. rugosus* Emery, *P. occidentalis* Cresson, and *P. salinus* Olsen species tend to harvest near the trunk of their foraging trails which are shaped by seed distribution, disturbances, or inter/intra-species interactions (MacMahon, 2000; Traniello, 1989).

Harvester ants exhibit seed preferences based on a combination of relative seed abundance, size/shape, and nutritional content of the seeds (Penn and Crist, 2018; MacMahon, 2000; Taber 1999). For instance, *P. occidentalis* Cresson prefers to forage with high species fidelity in seed-dense patches, which can reduce local seed bank heterogeneity (Luna et al. 2018; MacMahon, 2000; Crist and MacMahon, 1991). When the seed bank has low seed diversity, ants will collect less preferred seed varieties until more desirable seeds are available (MacMahon, 2000). When more preferred seeds return, ants will empty colony seed stores of the less desired seeds to replace them with preferred options (MacMahon, 2000).

Harvester ant seed foraging is not limited to natural areas and may occur in agricultural matrices where seed preferences may benefit or harm crop production. Although harvester ants are known to consume weed seeds, their seed preferences may also include consumption of crop seeds (Barbercheck and Wallace, 2021; Baraibar et al. 2011; Taber, 1999). Ant removal of crop seeds and vegetation causes economic loss, especially if the crop is situated within areas of high colony density (Reed and Landolt, 2019; Borth, 1982). Red harvester ants in particular are found in agricultural areas in the LRGV and may have a large impact on the plant community through removal of vegetation surrounding their nest entrance (1-5 m in diameter) or through seed collection (Reed and Landolt, 2019; MacMahon and Crist, 2000)

In addition to cash crops, harvester ants in agricultural fields may forage on cover crop seeds (based on personal communications) but have not been well documented. In the LRGV, cover crops are used during fallow periods to prevent soil erosion from wind or water (Soti and Racelis, 2020; Martinez et al., 2020; Nicolas Labrière, 2015; Bodner et al., 2010 Yu et al., 2000). Presumably, if an ant-preferred seed is sown within the foraging range of a colony, it will not have time to germinate before being taken by a forager to the colony granary. The lack of a root system and above-ground vegetation in the foraged field area can then potentially increase economic loss for the farmer (Soti and Racelis, 2020; Martinez et al., 2020; Bodner et al., 2010). So, preventing harvester ant interference with cover crops could potentially reduce soil exposure to erosion as well as save the cost of having to re-seed foraged areas.

The primary objective of the study was to determine red harvester ant preferences for commonly used cover crop seeds in the LRGV. We chose members of the families Fabaceae, Poaceae, and Brassicaceae that are currently being evaluated by farmers in the LRGV - hairy vetch (*Vicia villosa*), oat (*Avena sativa*), sunn hemp (*Crotalaria juncea*), radish (*Raphanus sativus*), and fescue (*Festuca arundinacea*). Wheatgrass (Poaceae: *Triticum aestivum*) was also included as a known preferred food for harvester ants and served as a control for the experiments (Brito-Bersi et al., 2018; Ryti and Case, 1988). Based on known baseline preferences that harvester ants exhibit towards grasses, we anticipated that oat, fescue, and wheatgrass would be most preferred as they are sugar-rich grasses from the family Poaceae. (MacMahon, 2000; Taber 1999). In addition to the use of cover crops, LRGV farmers may inoculate cover crop seeds with nitrogen-fixing bacteria to facilitate root nodulation to further benefit the soil (Rai et al., 2021; Kasper et al., 2019; Kasper 2019). As such treatments may influence ant foraging decisions, the second objective was to determine if seed inoculation treatments used for increased germination rates would alter the previously established cover crop seed preferences.

## Methods

### Site Description

The study site was located within the Lower Rio Grande Valley in South Texas. This area is considered a local steppe climate that is subtropical subhumid marine with an average annual temperature of 24°C (16.3-30.2°C) and 572 mm of precipitation. Soils in these regions of the Rio Grande Plain are considered deep loamy soils with moderately sloped planes and an average altitude of 34 m (USDA, 2008). Specifically, all trials were conducted at the University of Texas at Rio Grande Valley (UTRGV) campus (~ 1.5 km^2^) in Edinburg, Hidalgo County, TX, USA (26.306667, −98.170944). This site was selected as the ants present would have no prior exposure to the species of seeds presented during the study, but would also still experience disturbance pressures such as irrigation and routine mowing (a proxy for agricultural practices relative to natural settings).

On the campus, most vegetation included grasses used for lawns intermixed with weeds (primarily grass burr/sticker burr) and punctuated by standard suburban ornamental plants (such as Tropical Milkweed (*Asclepias curassavica*) and *Lantana sp.*). As of publication of the 2020 Tree Campus USA Report, there are 53 different species of trees with Live Oak (*Quercus virginiana*), Texas Ebony (*Ebenopsis ebano*), and Honey Mesquite (*Prosopis glandulosa*) being noteworthy examples (UTRGV Office For Sustainability, 2021). The immediate land use surrounding the study site is considered a combination of suburban and peri-urban with intermixed sorghum fields, pasture, and citrus groves. Land use within the LRGV more generally also includes mixed fruit and vegetable crops as well as sugarcane production. Active *Pogonomyrmex* colonies (n = 37) with no prior exposure to cover crop seeds near were mapped throughout the site using an eXplorist 610 GPS unit (Magellan, San Dimas, CA, USA). Colony activity was determined by whether there were foraging trails present with active bidirectional ant traffic.

### Seed Preference Trials

To determine whether size differences between seeds could impact preference, 10 seeds of each variety were weighed and averaged and seed texture was noted. For the trial, the cover crop seeds - hairy vetch (Johnny’s Selected Seeds, Winslow, ME), oat, sunn hemp (Johnny’s Selected Seeds, Winslow, ME), wheatgrass (Todd’s Seeds, Livonia, MI), radish (Johnny’s Selected Seeds, Winslow, ME), and fescue (GreenCover, Bladen, NE) - were pre-counted in groups of 10 seeds per cover crop and stored in microcentrifuge tubes at room temperature before transport to the field. Seed depots were constructed out of I-plate Petri dishes (100 mm × 15 mm). Petri dishes were sanded to produce a rough surface to increase traction, and 3 U-shaped entrances were created with a soldering iron at 45° and 90° angles on each half of the Petri dish to allow for easy ant entry to the dish.

The seed depot was placed 2 m from the nest entrance along the primary foraging trail with seed depot entrances facing the foraging trail (see supplemental materials for optimization of depot placement and depot construction). Upon initiation of each trial, the seeds were placed into a depot, with even numbers per side and a total of 10 seeds per cover crop available per colony. After the addition of the seeds, cages (1 cm × 1 cm hardware cloth [Everbilt, The Home Depot, Atlanta, GA] shaped into a 23 cm × 23 cm square) were placed on top of the depots and secured into the ground with 3 cm fence staples to prevent vertebrate removal of the seeds and indicate human interference (Campagnoli and Christianini, 2021; Thompson et al., 2016; Hughes and Westoby, 1990). Seed removal was documented at intervals of 1, 2, 4, and 24 h. During each inspection, temperature, wind speed, and cloud cover percentage were measured and the seeds within and outside of the depots were counted. Seed preference trials were conducted from February to June 2020 in groups of 8-10 colonies per observation period. The tested colonies (n=37) were a minimum of 10 m apart to prevent overlap of colony foraging. All trials were conducted within a temperature range of 20.5-36.6°C and wind speeds ≤ 32km/h to optimize ant foraging time but minimize the risk of wind overturning the seed depots.

Due to a delay in shipping, the colonies observed in the first two days of trials (n=12) were not immediately exposed to fescue seeds. These colonies were re-tested later with a depot mix including fescue seeds. They were compared to colonies that were exposed to fescue from the start and they did not demonstrate any difference in preference. Because of this lack of difference, we decided to use the full data set from the second round of trials from the initial twelve colonies for data analyses.

### Seed Inoculation Trials

The experimental design for the seed inoculation trials was conducted in a similar manner to the seed preference trials. The same colonies (n=34) and number of colonies per observation period (n=8-10) were used. To differentiate which side held inoculated seeds and which held non-inoculated, the underside of depots were marked with a small section of tape. Two preferred seeds from the seed preference trials belonging to different plant families (wheatgrass and radish) were used to ensure that any inoculation effects would not be confused with lack of preference. Seeds were inoculated in the laboratory with the Guard-N Omri Seed Inoculant (Johnny’s Selected Seeds, Winslow, ME) via slurry method. For every 90 g of seeds, 0.7 g of inoculant was added to the container and shaken. Seeds were stored at room temperature in a marked microcentrifuge tube until use in the field. Trials were completed between July and August 2020 according to the previously used seed preference methods.

### Statistical Analysis

R version 3.6.2 (RStudio Team, 2020) was used to conduct all statistical analyses. Within each dataset, each seeds’ time to removal was categorized individually with censoring due to external events (e.g., flipped depots due to high wind speeds, removal of the cage prior to the 24 hours period, etc.) denoted. The survdiff function from the survival package was used to determine if there was a significant difference in ant cover crop preference (Therneau, 2015; Therneau and Grambsch, 2000).The Kaplan-Meier survival estimator, which estimates the likelihood of an event occurring at a point in time, was used to calculate seed removal event likelihood over time (Johnson, 2018). The log-rank test using the lifelines package (Rickert, 2017), a hypothesis test that compares the survival distribution between two samples, was used to compare the survival distribution of cover crop seeds to the wheatgrass and non-inoculated controls. To further investigate these differences while incorporating other variables such as observation month, we used Cox proportional hazard models and preferences compared against the wheatgrass standard using the ggforest function from the survival package (Therneau, 2015; Therneau and Grambsch, 2000).

## Results

### Seed Preference Trials

Kaplan Meier survival curves were used to compare removal rates of the different cover crop seed varieties (Fig. 1). The Cox proportional hazards model determined the only significant differences in removal were between wheatgrass and vetch (*p* < 0.001), wheatgrass and sunn hemp (*p* < 0.001), and wheatgrass and fescue (*p* < 0.050), (Fig. 2; Table 1). During the trials, ants exhibited a preference for wheatgrass and oat seeds, often removing all the seeds before 24 h (Table 1). For differences between seed types outside of wheatgrass, a pairwise log rank test was used.

**Figure 1.**
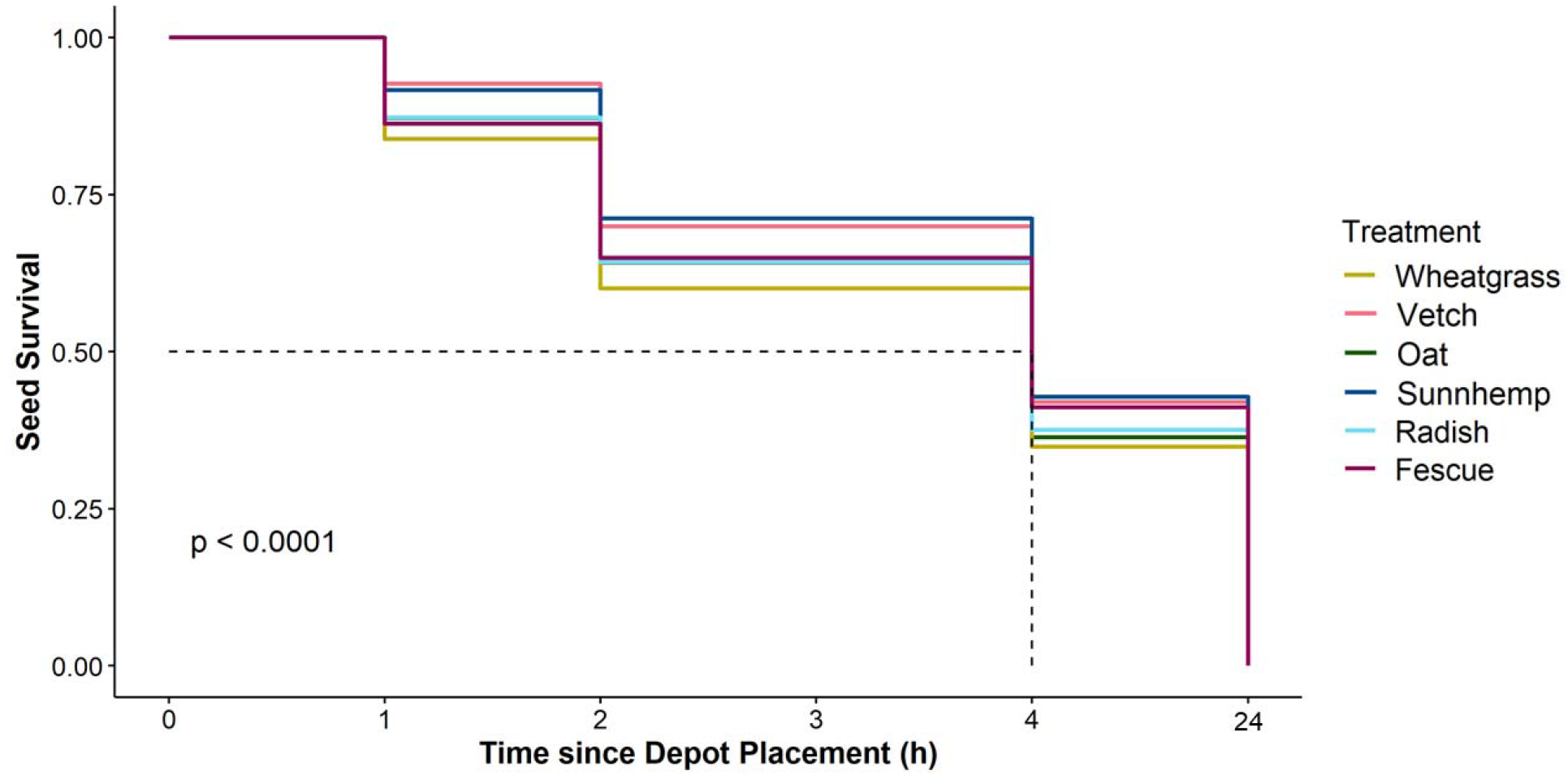
Kaplan Meier curve of seed types’ likelihood of survival over the course of the seed preference trial based on selected data. (n=37 colonies). The dashed line indicates the overall median removal time.

**Figure 2.**
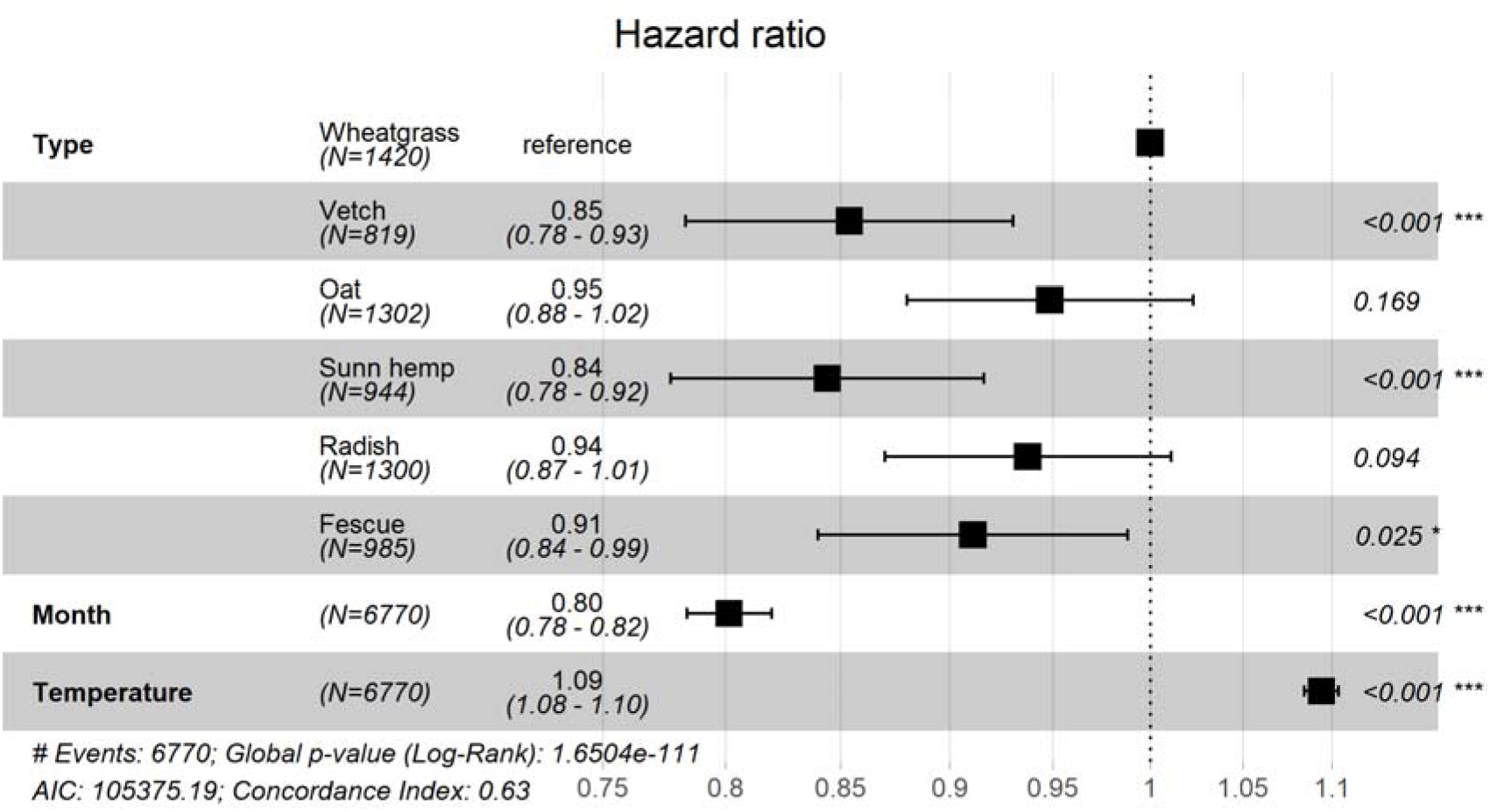
Hazard Proportional Ratio test demonstrating differences in preferences between seed types. Reference is wheatgrass. Means on the right side of the chart indicate a larger number of seeds that were removed during the trial. Differences in n (observed seed number) were due to seeds that were censored for external events.

**Table 1.**
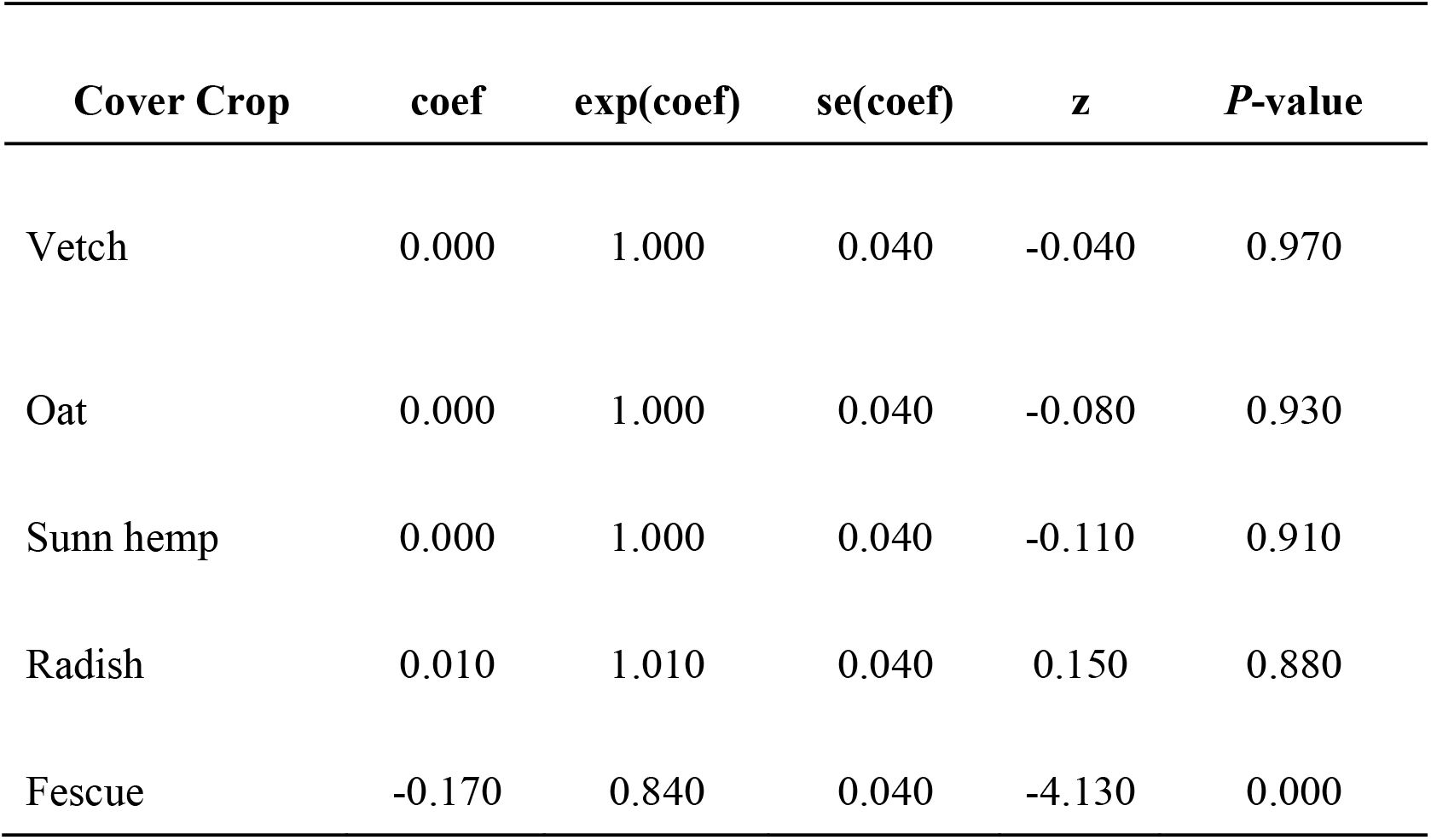
Summary of the fitted cox model for cover crop seed preferences.

The pairwise log rank test provided differences in survival between the seeds amongst themselves (Table 2). Vetch and Sunnhemp, though not significantly different from one another, were the varieties that were significantly less harvested in comparison to other seed types outside of wheatgrass. Overall, vetch was found to be significantly less harvested when compared to oat (p < 0.050), wheatgrass (p < 0.001) or radish (p < 0.050) Sunn hemp was found to be significantly less harvested when compared to oat (p < 0.005), wheatgrass (p < 0.001), or radish (p < 0.003) (Fig. 2; Table 2). Similarly to the Cox proportional hazards model, Fescue, while not being significantly different from vetch or sunn hemp, was significantly less harvested than wheatgrass (p < 0.050), another member of the Poaceae family. Other than seed types, seed collection differed among months (Supplementary Fig. 1). Over time, seed collection significantly decreased from February to June (Supplementary Fig. 1).

**Table 2.** Pairwise comparisons using Log-Rank Test between seed types for the seed preference study (n = 6770 total seeds). Levels of significance indicated by asterisks.

|  | **Vetch** | **Oat** | **Sunn hemp** | **Wheatgrass** | **Radish** |
| --- | --- | --- | --- | --- | --- |
| **Oat** | 0.003 |  |  |  |  |
| **Sunn hemp** | 0.733 | <0.001 |  |  |  |
| **Wheatgrass** | <0.001 | 0.149 | <0.001 |  |  |
| **Radish** | 0.011 | 0.664 | 0.003 | 0.068 |  |
| **Fescue** | 0.190 | 0.121 | 0.115 | 0.003 | 0.230 |

The physical characteristics of the seeds in the depot did not appear to affect preference as the preferred seeds in the study did not consistently share characteristics. Non-prefered seeds also did not share seed shape or texture, only color and nitrogen-fixing abilities. All the seeds’ weights were similar with the exception of fescue and radish, which were significantly lighter than the other varieties (Fig. 3). Vetch and radish shared physical characteristics-both were round and uneven in texture, but they were treated differently by the ants. Sunn hemp was smooth, and bean shaped, while oat and fescue appeared fibrous towards the ends with a thin and elongated shape. Wheatgrass was oblong in shape and relatively smooth.

**Figure 3.**
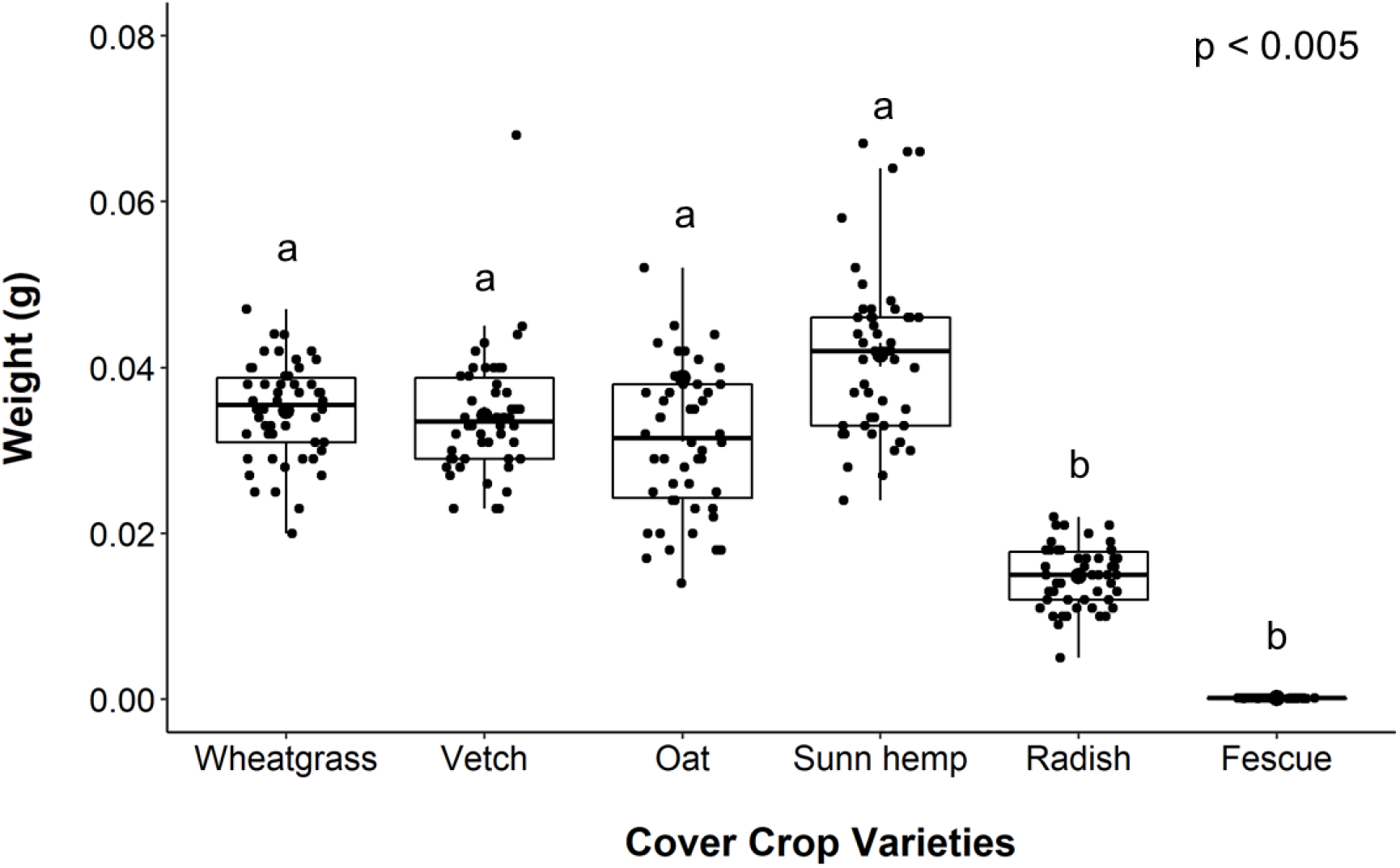
Differences in seed weight between the six cover crop seeds (n=50/seed type) used in the study. Boxplots are in the style of Tukey where the box limits represent the lower 25% and upper 75% quantile with the line representing the median. Tukey HSD was used to determine significance differences (denoted by letters) among seed weights.

### Seed Inoculum Trial

Unlike the seed preference trials, inoculum trials did not indicate significant differences in preference. The Kaplan Meier curve created from the collected data further demonstrated the visual lack of preference between inoculated versus non-inoculated seed between the same seed type (Fig. 4). Additionally, the Cox proportional hazards data demonstrated that the difference in preference between the inoculated and non-inoculated seeds was not significant (Fig. 5; Table 3). This lack of overall preference also meant that there was no preference between one another (Table 4). Surprisingly, the only significance found within the trial was a change in seed removal (Supplementary Fig. 2). Depot harvesting was significantly higher in July in comparison to June or August.

**Figure 4.**
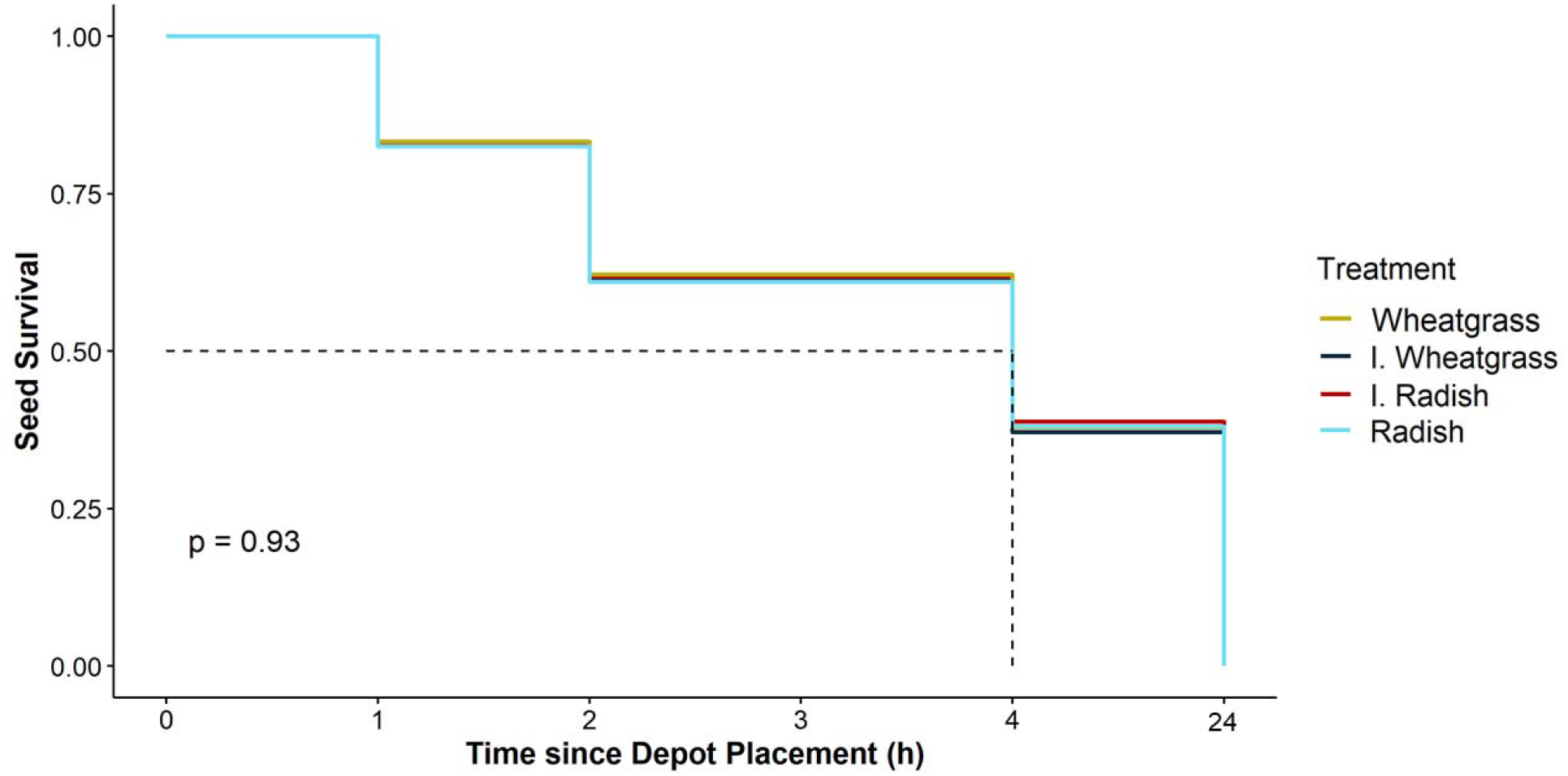
Kaplan Meier curve of seed types’ likelihood of survival over the course of the seed inoculation trial based on selected data. (n=34 colonies). The dashed line indicates the overall median removal time.

**Figure 5.**
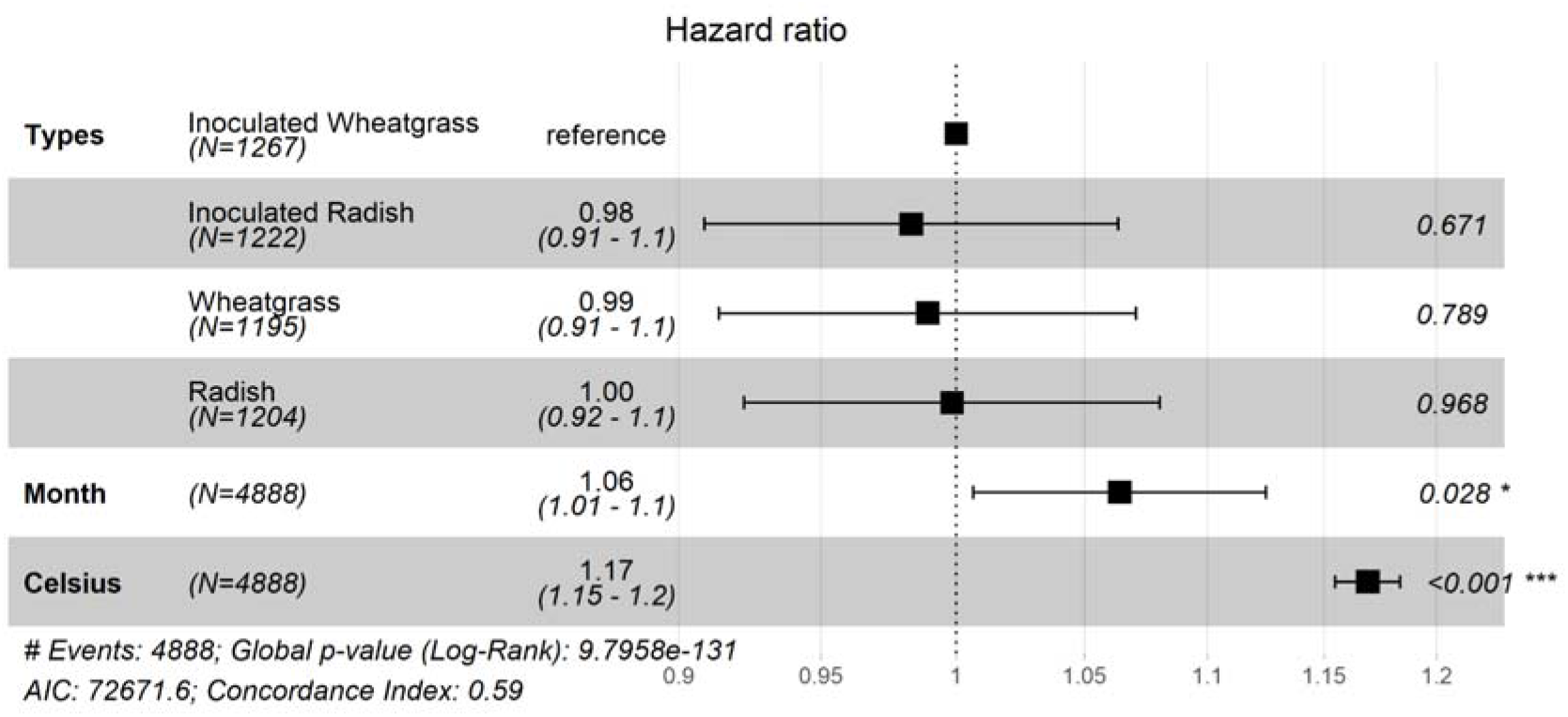
Hazard Proportional Ratio test demonstrating differences in preferences between inoculated and uninoculated seed types. Reference is wheatgrass. Means on the right side of the chart indicate a larger number of seeds that were removed during the trial. Differences in n (observed seed number) were censored for external events.

**Table 3.** Summary of the fitted cox model for inoculated seed preferences.

| Cover Crop | coef | exp(coef) | se(coef) | z | Pr(>\|z\|) |
| --- | --- | --- | --- | --- | --- |
| Inoculated Wheatgrass | 0.010 | 1.010 | 0.040 | 0.260 | 0.750 |
| Inoculated Radish | -0.010 | 0.990 | 0.040 | 0.680 | 0.820 |
| Radish | 0.000 | 1.010 | 0.040 | 0.410 | 0.930 |

## Discussion

The goal of the study was to determine if red harvester ants exhibit preferences among different cover crop seed varieties and whether inoculating preferred seed types with nitrogen-fixing bacteria would inhibit the desirability of the seed. We introduced naive harvester ants to agricultural seeds via seed depots deployed over 24 h. We found that harvester ants had a significant preference for grass seeds and radish seeds compared to nitrogen-fixing sunn hemp and vetch seeds. However, we did not observe any difference in preference between inoculated and non-inoculated seeds of either wheatgrass or radish.

We had assumed harvester ants would prefer to forage on certain seeds based on physical characteristics and family (Poaceae) (Penn and Crist, 2018; MacMahon, 2000; Taber 1999). As anticipated due to prior work on seed preferences in natural areas, all grass seeds were similarly preferred. However, the attributes of radish overlapped with the less preferred seeds in terms of shape, color, or weight, indicating these physical trails were not the only driver of preference within this context (MacMahon, 2000; Taber 1999).

Alternatively, seed preferences could have been based on seed availability in the surrounding habitat, which. likely changed from February 2020 to August 2020. During the study, we observed native seed burrs (Genus *Cenchrus* L.) being taken into the colony often as well as smaller grass seeds. Prior documentation of burrs in and around Hidalgo county indicates that burrs are annual grasses with an affinity for frequently disturbed sites such as roadsides, similar to the study sites (Goel et al., 2011; Shaw, 2011). *Cenchrus echinatus* L. begin to germinate in the late spring, continuing through the fall (Smith et al., 2012; Cope & Gray, 2009). The decrease in seed removal from trials that occurred from spring to summer could be a change in priority from depot seeds to collecting recently germinated seeds from the surrounding *Cenchrus sp*. Given these observations, the interactions of cover crop seeds with weed banks within agricultural settings needs to be evaluated further, particularly in regards to sowing timing. Outside of seed preference changes due to the surrounding seed pool, *P. barbatus* activity is closely related to rainfall, peaking in the summer months and correlated with overall seed availability. With additional rainfall, more grasses outside of drought resistant varieties such as *Cenchrus sp*. potentially germinated, allowing for more diversity in the seed pool (MacMahon, 2000; Smith et al., 2012; Cope & Gray, 2009). The additional surrounding native seeds could have been another cause for the reduction in depot harvesting over time from February to June. Alternatively, during the sudden increase in depot harvesting from June to July could be in preparation for August, which is usually known for its higher temperatures. In August, activity significantly decreased in comparison to both June and July, implying that high amounts of collection in June could have been done to avoid excess water loss for the colonies in August (Supplementary Fig. 2)

Another interesting, isolated event was recorded on July 23rd, 2020, two days prior to the touch down of Hurricane Hanna in the LRGV. Within one hour, 8 of 9 colonies had completely emptied the depots. The impacts of such weather events are known to affect insect behavior in response to changes in barometric pressure; many insects exhibit sudden insatiable appetites likely preparing for weather events that follow. (Fernando R. Sujimoto, 2019; Flitters, 1963). Leaf-cutter ants have been observed to significantly increase foraging during periods of low barometric pressure, and harvester ants may do the same (Fernando R. Sujimoto, 2019). Future studies regarding the correlation between harvester ant foraging intensity and barometric pressure could help determine risk during certain planting dates in regions along the gulf coast that have the potential to experience tropical cyclones annually.

Harvester ants have been previously observed to have contradictory behavior regarding the same seed species based on other aspects such as seed germination or fungal infection (MacMahon et al., 2000; Taber, 1999; Crist and Friese, 1993). However, in our trials, inoculated and non-inoculated seeds were not treated differently, indicating that the presence of nitrogen-fixing bacteria did not inhibit or encourage harvester ant predation. Regardless, there is conflicting data regarding the amount of microbial diversity/biomass within the soil around ant colonies (Ginzburg et al., 2008; Boulton et al., 2003; Wagner et al., 1997). *Pogonomyrmex barbatus* in the study showed no preference towards or against inoculated seeds, hinting that their granaries could be potentially rich in microbial activity. Alternatively, harvester ants do partake in seed cleaning behavior that could occur at any point prior to introduction to the granary.

In subtropical areas such as the LRGV, prior studies recommend the use of warm season cover crops due to subtropical climate and promotion of native mycorrhizal fungi (Soti et al. 2016; Rugg, 2016). Based on the data collected in this study, harvester ants were exhibited lower levels of preference towards certain seed varieties such as sunn hemp. The benefits that these nitrogen fixing varieties, such as sunn hemp, hold towards the soil can be extremely beneficial. Sunn hemp, for example, conserves phosphorus in the soil, increases nitrates, and has the potential to improve soil health in subtropical agroecosystems such as the LRGV (Soti et al. 2016; Rugg, 2016; Mansoar et al. 1997). Not only does sunn hemp have the potential to be an excellent South Texas cover crop, but it is also increasing in popularity in other southern areas of the U.S. like Florida and Louisiana. Similarly, hairy vetch also has potential to be a great cover crop due to the low ant preference and its weed suppression and nitrogen-fixing abilities (Moran and Greenberg, 2008). Given these cover crops are not preferred over grasses in the seed depot study, which are common in the non-crop habitats surrounding LRGV crop fields, harvester ants would likely predate on surrounding weeds and grasses instead of the chosen cover crop.

Harvester ants can be a substantial disturbance agent in arid to semi-arid regions of the United States and Mexico. *Pogonomyrmex sp*. have a pest status for seed collection and plant removal in agricultural areas and can remove up to 100% of a preferred seed within their foraging range (Crist and MacMahon, 1992; Tabber, 1999). Our data suggests we can recommend nitrogen-fixing cover crops like sunn hemp and vetch to farmers as a potential cover crop during fallow periods and could be paired with the fact that seed inoculation is neither preferred or rejected by harvester ants. Inoculating these nitrogen-fixing seeds could help with nodulation, nitrogen-fixing processes, and benefit the soil health below ground while protecting topsoil from erosion. Not only that, using the pair for a cover crop trial, could in turn encourage harvester ant predation on weed species or surrounding native plants that could limit crop yields (Baraibar et al., 2011). Additional research should be conducted regarding harvester ant preferences. For example, conducting preference studies with rural harvester ants that have more exposure to different agricultural seed varieties and in turn, potential differences in preferences. A better understanding of harvester ant seed preferences can be used to encourage predation on native or weed seeds while reducing the need to eradicate native harvester ant colonies.

## Acknowledgements

Funding support for LE was provided by the Dean’s Graduate Research Assistantship from UTRGV. We would like to thank the Racelis’ Agroecology Lab at UTRGV for providing the seeds used in the experiment, as well as the inoculum.

## Literature Cited

Baraibar, B., Ledesma, R., Royo-Esnal, A., & Westerman, P. R. (2011). Assessing yield losses caused by the harvester ant *Messor barbarus* (L.) in winter cereals. Crop Protection, 30(9), 1144–1148.

Barbercheck, M. E., & Wallace, J. (2021). Weed–Insect Interactions in Annual Cropping Systems. Annals of the Entomological Society of America, 114(2), 276–291.

Bodner, G., Himmelbauer, M., Loiskandl, W., & Kaul, H. P. (2010). Improved evaluation of cover crop species by growth and root factors. Agronomy for sustainable development, 30(2), 455–464.

Borth, P. W. T., B. R.; Johnson, G. D. (1982). A Preliminary Evaluation of Amdro for Control of a Harvester Ant *(Pogonomyrmex maricopa* Wheeler) in Hard Red Spring Wheat. Forage and Grain: A College of Agriculture Report. 39–41.

Boulton, A. M., Jaffee, B. A., & Scow, K. M. (2003). Effects of a common harvester ant *(Messor andrei)* on richness and abundance of soil biota. Applied Soil Ecology, 23(3), 257–265.

Brito-Bersi, T., Dawes, E., Martinez, R., & McDonald, A. (2018). Seed preference in a desert harvester ant, *Messor pergandei*. California Ecology and Conservation Research, 1–6.

Campagnoli, M. L., & Christianini, A. V. (2021). Temporal consistency in interactions among birds, ants, and plants in a neotropical savanna. Nordic Society Oikos, 00: 1–13

Cope, T. & Gray, A. (2009). Grasses of the British Isles. Botanical Society of the British Isles No.13.

Crist, T. O., & Friese, C. F. (1993). The impact of fungi on soil seeds: implications for plants and granivores in a semiarid shrub□steppe. Ecology, 74(8), 2231–2239.

Crist, T. O., & MacMahon, J. A. (1992). Harvester ant foraging and shrub□steppe seeds: interactions of seed resources and seed use. Ecology, 73(5), 1768–1779.

Davidson, D. W. (1982). Sexual selection in harvester ants (Hymenoptera: Formicidae: Pogonomyrmex). Behavioral Ecology and Sociobiology, 10(4), 245–250.

Davis, J. M. (2016). Management of the Red Harvester Ant *Pogonomyrmex barbatus*. Texas Parks and Wildlife Department. https://tpwd.texas.gov/huntwild/wild/wildlife_diversity/texas_nature_trackers/horned_lizard/documents/harvester_ant_management.pdf

Flitters, N. E. (1963). Observations on the effect of hurricane “Carla” on insect activity. International Journal of Biometeorology, 6(2), 85–90.

Friese, C. F., & Allen, M. F. (1993). The interaction of harvester ants and vesicular-arbuscular mycorrhizal fungi in a patchy semi-arid environment: the effects of mound structure on fungal dispersion and establishment. Functional Ecology, 13–20.

Ginzburg, O., Whitford, W. G., & Steinberger, Y. (2008). Effects of harvester ant *(Messor spp*.) activity on soil properties and microbial communities in a Negev Desert ecosystem. Biology and fertility of Soils, 45(2), 165–173.

Goel S., Singh H.D., Raina S.N. (2011) Cenchrus. In: Kole C. (eds) Wild Crop Relatives: Genomic and Breeding Resources. Springer, Berlin, Heidelberg. https://doi.org/10.1007/978-3-642-14255-0_3

Graeber, K. N., and G Leubner-Metzger,. (2017). Encyclopedia of Applied Plant Sciences (Second Edition). 1, 483–489. doi: https://doi.org/10.1016/B978-0-12-394807-6.00209-4

Hughes, L., & Westoby, M. (1990). Removal Rates of Seeds Adapted for Dispersal by Ants. Ecology, 71(1), 138–148. doi: https://doi.org/10.2307/1940254

Johnson, L. L. (2018). An Introduction to Survival Analysis. Principles and Practice of Clinical Research, 4, 373–381. doi:10.1016/B978-0-12-849905-4.00026-5

Johnson, R. A. (1998). Foundress survival and brood production in the desert seed-harvester ants *Pogonomyrmex rugosus* and *P. barbatus* (Hymenoptera, Formicidae). Insectes Sociaux, 45(3), 255–266.

Laundré, J. W. (1990). Soil moisture patterns below mounds of harvester ants. Rangeland Ecology & Management/Journal of Range Management Archives, 43(1), 10–12.

Kasper, S. L. (2019). Investigating Limitations to Nitrogen Fixation by Leguminous Cover Crops in South Texas. The University of Texas Rio Grande Valley.

Kasper, S., Christoffersen, B., Soti, P., & Racelis, A. (2019). Abiotic and biotic limitations to nodulation by leguminous cover crops in South Texas. Agriculture, 9(10), 209.

MacKay, W. P., & Mackay, E. (2002). The ants of New Mexico (Hymenoptera: Formicidae) (p. 400). Lewiston, NY: Edwin Mellen Press.

MacMahon, J. A., Mull, J. F., & Crist, T. O. (2000). Harvester ants *(Pogonomyrmex spp.):* their community and ecosystem influences. Annual Review of Ecology and Systematics, 31(1), 265–291.

Mansoer, Z., Reeves, D. W., & Wood, C. W. (1997). Suitability of sunn hemp as an alternative late summer legume cover crop. Soil Science Society of America Journal, 61(1), 246–253.

Moran, P. J., & Greenberg, S. M. (2008). Winter cover crops and vinegar for early-season weed control in sustainable cotton. Journal of Sustainable Agriculture, 32(3), 483–506.

Labrière, N., Locatelli, B., Laumonier, Y., Freycon, V., & Bernoux, M. (2015). Soil erosion in the humid tropics: A systematic quantitative review. Agriculture, Ecosystems & Environment, 203, 127–139.

Penn, H. J., & Crist, T. O. (2018). From dispersal to predation: A global synthesis of ant–seed interactions. Ecology and evolution, 8(18), 9122–9138.

Rai, Q., Choudhury, R. A., Soti, P., & Racelis, A. (2021). Rhizobial adhesives enhance nodule formation in sunn hemp. bioRxiv.

Reed, H. C., & Landolt, P. J. (2019). Ants, wasps, and bees (Hymenoptera). In Medical and veterinary entomology (pp. 459–488). Academic Press.

Rickert, J. (2017). Survival analysis with R. In: RStudio (Ed.) R Views: R Community Blog. Boston, MA. Retrieved from https://rviews.rstudio.com/2017/09/25/survival-analysis-with-r/.

RStudio Team (2020). RStudio: Integrated Development for R. RStudio, PBC, Boston, MA URL http://www.rstudio.com/.

Rugg, S. M. (2016). Multifunctionality of Cover Crops on Organic Vegetable Farms in South Texas. The University of Texas Rio Grande Valley.

Ryti, R. T., & Case, T. J. (1988). Field experiments on desert ants: testing for competition between colonies. Ecology, 69(6), 1993–2003.

Shakeel, M., Ali, H., Ahmad, S., Said, F., Khan, K. A., Bashir, M. A.,… Ali, H. (2019). Insect pollinators diversity and abundance in *Eruca sativa Mill.* (Arugula) and *Brassica rapa L*. (Field mustard) crops. Saudi Journal of Biological Sciences, 26(7), 1704–1709. doi: https://doi.org/10.1016/j.sjbs.2018.08.012.

Shaw, R. B. (2011). Guide to Texas grasses. Texas A&M University Press.

Smith, H., Ferrell, J., & Sellers, B. (2012). Identification and Control of Southern Sandbur *(Cenchrus echinatus* L.) in Hayfields. EDIS, 2012(12).

Snyder, S. R., & Friese, C. F. (2001). A survey of arbuscular mycorrhizal fungal root inoculum associated with harvester ant nests (*Pogonomyrmex occidentalis*) across the western United States. Mycorrhiza, 11(3), 163–165.

Soti, P., & Racelis, A. (2020). Cover crops for weed suppression in organic vegetable systems in semiarid subtropical Texas. Organic Agriculture, 10(4), 429–436.

Sujimoto, C. M. C., Caio H. L. Zitelli, José Maurício S. Bento. (2019). Foraging activity of leaf cutter ants is affected by barometric pressure. Ethology. doi:10.1111/eth.12967.

Terry M. Therneau, Patricia M. Grambsch (2000). _Modeling Survival Data: Extending the Cox Model_. Springer, New York. ISBN 0-387-98784-3.

Therneau T (2015). _A Package for Survival Analysis in S_. version 2.38, <URL:https://CRAN.R-project.org/package=survival>.

Thomson, F. J., Auld, T. D., Ramp, D., & Kingsford, R. T. (2016). A switch in keystone seed-dispersing ant genera between two elevations for a myrmecochorous plant, Acacia terminalis. Plos one, 11(6), e0157632.

Tizon, F. R., Peláez, D. V., & Elía, O. R. (2010). Efecto de los cortafuegos sobre el ensamble de hormigas (Hymenoptera, Formicidae) en una región semiárida, Argentina. Iheringia. Série Zoología, 100, 216–221.

Treadwell, D. D., & Alligood, M. (2008). Sunn hemp (*Crotalaria juncea L*.): A summer cover crop for Florida vegetable producers. EDIS, 2008(2).

UTRGV Office For Sustainability. (2021). 2020 Tree Campus USA Report. Issuu. Retrieved December 2, 2021, from https://issuu.com/officeforsustainability/docs/2020_tree_campus_report-_final_5_.

Vieira□Neto, E. H., Vasconcelos, H. L., & Bruna, E. M. (2016). Roads increase population growth rates of a native leaf□cutter ant in Neotropical savannas. Journal of Applied Ecology, 53(4), 983–992.

Uhey, D. A., Cummins, G. C., Rotter, M. C., Lassiter, L. S., & Whitham, T. G. (2021). Hiking Trails Increase Abundance of Harvester Ant1 Nests at Clear Creek, Arizona. Southwestern Entomologist, 46(2), 403–412.

United States Department of Agriculture Natural Resources Conservation Service. (2008). Texas General Soil Map. General Soil Map of Texas. Retrieved November 19, 2021, from https://maps.lib.utexas.edu/maps/texas/texas-general_soil_map-2008.pdf.

