## Supplementary Materials for "Red harvester ant (Order: Hymenoptera) preference for cover crop seeds in South Texas"

### Supplementary Methodology

#### *Depot Construction*

Sections of hardware cloth were cut in 4 sections of 7 cm x 23 cm pieces and 1 23 cm x 23 cm cube.  The ends of the wire were tripped neat to prevent injury to members of the lab. The 4 7 cm x 23 cm sections were tied together at the short ends to one another using zip-ties to make the outer form of the box. Excess ends of the zip-tie were trimmed off to facilitate transport. When the four sections were secure their long ends were secured to the perimeter of the  23 cm x 23 cm base using zip-ties.

#### *Optimization of depot placement*

Seeds were placed ½ m, 1m, 2m, and 3m away from the colony to monitor ant interactions with the depots.  Visual observations were made to see when seeds were most often noticed by foraging harvester ants. When the depots were located too close to the colony ( ½ m - 1m away), other members of the colony such as members of the waste disposal unit would interfere or be disrupted by the depot in their path. Alternatively, depots located too far (3m) had less interactions with foragers as the foraging trail had begun to branch off into smaller section and foragers were more widely dispersed. Placing the depot 2m away from the colony allowed for higher concentrations of foraging ants to interact with the depot on their way from and to the colony.

**Figures**

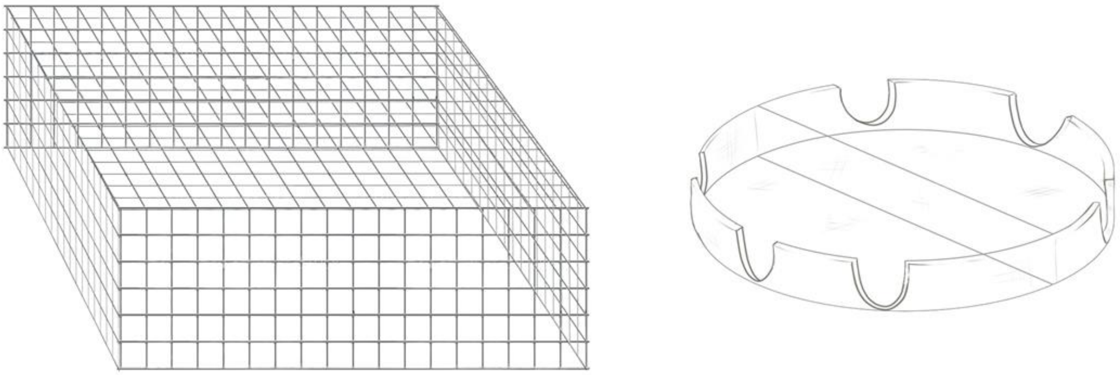

a) b)

**Supplementary Diagram 1.** Diagrams of the depot constructed from wire to protect seeds from larger herbivores such as birds and rodents (left) and the modified I-plated petri dish for seed holding (right).

Figure 1

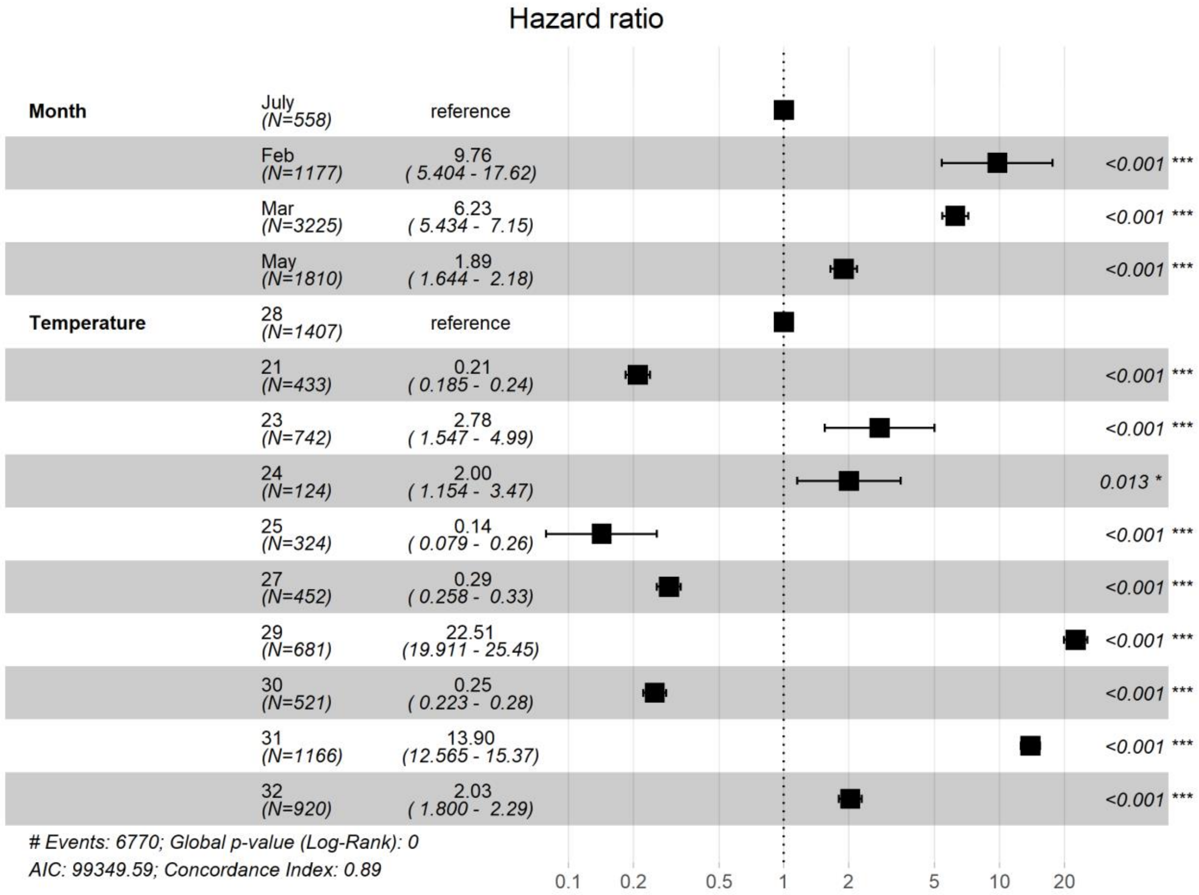

**Supplementary Figure 1.** Hazard Proportional Ratio test demonstrating differences in cover crop seed preferences between seeds taken during the course of the experiment at different months and temperatures. Differences in Month n (observed seed number) was differences in numbers of trials per month and different temperatures at the time of seed counts.

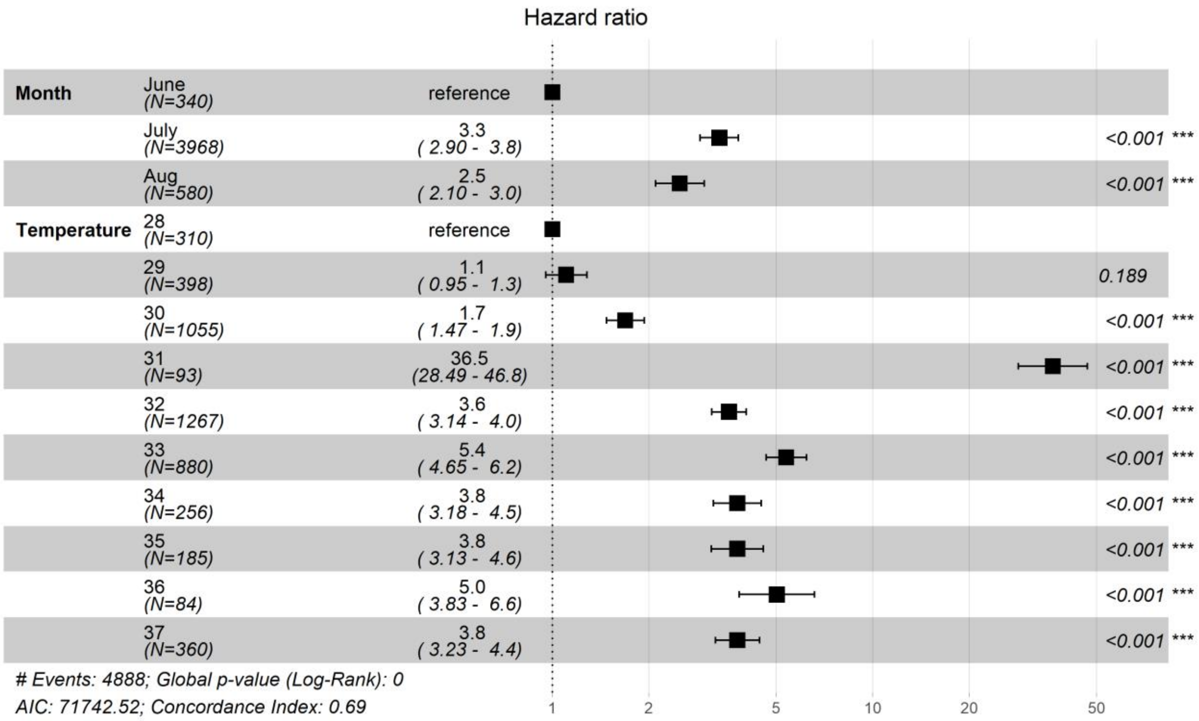

**Supplementary Figure 2.** Hazard Proportional Ratio test demonstrating differences in depot seed removal at varying months and temperatures over the course of the inoculation trials. Differences in Month n (observed seed number) was differences in numbers of trials per month and temperatures at the time of seed count.

**Supplementary Table 1**. Summary of the fitted cox model for cover crop seed preferences.

| Type | coef | Exp(coef) | Se(coef) | z | P r(>\|z\|) |
|---|---|---|---|---|---|
| Vetch | -0.149 | 0.860 | 0.043 | -3.411 | <0.001*** |

| | | | | | |
|---|---|---|---|---|---|
| Oat | -0.044 | 0.956 | 0.038 | -1.153 | 0.248 |
| Sunn Hemp | -0.161 | 0.850 | 0.042 | -3.847 | <0.001*** |
| Radish | -0.060 | 0.940 | 0.038 | -1.586 | 0.112 |
| Fescue | -0.107 | 0.898 | 0.041 | -2.581 | 0.009 |

| Month | coef | Exp(coef) | Se(coef) | z | P r(>\|z\|) |
|---|---|---|---|---|---|
| Feb | 2.277 | 9.756 | 0.301 | 7.556 | 4.15e-14*** |
| Mar | 1.829 | 6.230 | 0.069 | 26.191 | <2e-16*** |
| May | 0.638 | 1.893 | 0.071 | 8.876 | <2e-16*** |

| Temperature | coef | Exp(coef) | Se(coef) | Z | P r(>\|z\|) |
|---|---|---|---|---|---|
| 21 | -1.558 | 0.21042 | 0.065 | -23.662 | <2e-16*** |
| 23 | 1.022 | 2.779 | 0.065 | 3.421 | 0.000623*** |
| 24 | 0.694 | 2.001 | 0.280 | 2.471 | 0.013482* |
| 25 | -1.946 | 0.142 | 0.301 | -6.449 | 1.13e-10*** |
| 27 | -1.227 | 0.293 | 0.065 | -18.847 | <2e-16*** |
| 29 | 3.114 | 22.512 | 0.062 | 49.707 | <2e-16*** |
| 30 | -1.379 | 0.251 | 0.062 | -21.975 | <2e-16*** |
| 31 | 2.631 | 13.897 | 0.051 | 51.189 | <2e-16*** |
| 32 | 0.709 | 2.032 | 0.061 | 11.438 | <2e-16*** |

| Month | coef | Exp(coef) | Lower .95 | Upper .95 |
|---|---|---|---|---|
| Feb | 9.756 | 0.102 | 5.403 | 17.615 |
| Mar | 6.230 | 0.160 | 5.433 | 7.145 |
| May | 1.893 | 0.528 | 1.644 | 2.179 |

| Temperature | Coef | Exp(coef) | Lower .95 | Upper .95 |
|---|---|---|---|---|
| 21 | 0.210 | 4.752 | 0.184 | 0.239 |
| 23 | 2.779 | 0.359 | 1.547 | 4.991 |
| 24 | 2.001 | 0.499 | 1.154 | 3.471 |
| 25 | 0.142 | 7.006 | 0.078 | 0.257 |
| 27 | 0.293 | 3.411 | 0.258 | 0.333 |
| 29 | 22.512 | 0.044 | 19.911 | 25.453 |
| 30 | 0.251 | 3.972 | 0.222 | 0.284 |
| 31 | 13.897 | 0.071 | 12.564 | 15.370 |
| 32 | 2.032 | 0.492 | 1.799 | 2.2947 |

| | | | | |
|---|---|---|---|---|
| | *** | ** | * | . |

| Signif. codes | 0.0001 | 0.001 | 0.01 | 0.05 | 0.1 | 1 |
|---|---|---|---|---|---|---|

**Supplementary Table 3**. Summary of the cox model fit for inoculated seed preferences.

| Type | Coef | exp(coef) | se(coef) | z | P r(>\|z\|) |
|---|---|---|---|---|---|
| Inoc. Wheatgrass | -0.012 | 0.987 | 0.040 | -0.320 | 0.749 |
| Inoc. Radish | -0.022 | 0.978 | 0.040 | -0.551 | 0.582 |
| Radish | -0.009 | 0.990 | 0.040 | -0.228 | 0.820 |

| Type | exp(coef) | Exp(-coef) | Lower .95 | Upper .95 |
|---|---|---|---|---|
| Inoc. Wheatgrass | 0.978 | 1.022 | 0.904 | 1.058 |
| Inoc. Radish | 0.987 | 1.013 | 0.912 | 1.068 |
| Radish | 0.990 | 1.009 | 0.915 | 1.072 |

| Month | coef | exp(coef) | se(coef) | z | P r(>\|z\|) |
|---|---|---|---|---|---|
| July | 1.200 | 3.321 | 0.069 | 17.154 | < 2e-16 *** |
| Aug | 0.913 | 2.499 | 0.089 | 10.291 | <2e16*** |

| Temperature | coef | exp(coef) | se(coef) | z | P r(>\|z\|) |
|---|---|---|---|---|---|
| 29 | 0.099 | 1.104 | 0.075 | 1.313 | 0.189 |
| 30 | 0.524 | 1.689 | 0.069 | 7.560 | 4.04e-14*** |
| 31 | 3.597 | 36.517 | 0.126 | 28.401 | <2e-16*** |
| 32 | 1.269 | 3.559 | 0.063 | 19.926 | <2e-16*** |
| 33 | 1.681 | 5.374 | 0.074 | 22.625 | <2e-16*** |
| 34 | 1.330 | 3.781 | 0.087 | 15.173 | <2e-16*** |
| 35 | 1.330 | 3.781 | 0.095 | 13.880 | <2e-16*** |
| 36 | 1.614 | 5.025 | 0.138 | 11.691 | <2e-16*** |
| 37 | 1.330 | 3.781 | 0.080 | 16.427 | <2e-16*** |

| Month | exp(coef) | exp(-coef) | lower .95 | upper.95 |
|---|---|---|---|---|
| July | 3.322 | 0.30107 | 2.89 | 3.810 |
| Aug | 2.499 | 0.40011 | 2.0992 | 2.976 |

| Temperature | exp(coef) | exp(-coef) | lower .95 | upper.95 |
|---|---|---|---|---|
| 29 | 1.105 | 0.90531 | 0.9522 | 1.281 |

| | | | | |
|---|---|---|---|---|
| 30 | 1.689 | 0.59194 | 1.4746 | 1.935 |
| 31 | 36.517 | 0.02738 | 28.4886 | 46.809 |
| 32 | 3.559 | 0.28095 | 3.1415 | 4.033 |
| 33 | 5.375 | 0.18606 | 4.6460 | 6.217 |
| 34 | 3.781 | 0.26445 | 3.1845 | 4.490 |
| 35 | 3.781 | 0.26445 | 3.1339 | 4.563 |
| 36 | 5.026 | 0.0.19899 | 3.8338 | 6.588 |
| 37 | 3.781 | 0.26445 | 3.2265 | 4.432 |

| | *** | ** | * | . | | |
|---|---|---|---|---|---|---|
| Signif. codes | 0.0001 | 0.001 | 0.01 | 0.05 | 0.1 | 1 |
